# A limited concentration range of diaphorin, a polyketide produced by a bacterial symbiont of the Asian citrus psyllid, promotes the *in vitro* gene expression with bacterial ribosomes

**DOI:** 10.1101/2024.01.05.574368

**Authors:** Rena Takasu, Takashi Izu, Atsushi Nakabachi

## Abstract

Diaphorin is a polyketide produced by “*Candidatus* Profftella armatura” (*Gammaproteobacteria*: *Burkholderiales*), an obligate symbiont of a devastating agricultural pest, the Asian citrus psyllid *Diaphorina citri* (Hemiptera: Psyllidae). Physiological concentrations of diaphorin, which *D. citri* contains at levels as high as 2–20 mM, are inhibitory to various eukaryotes and *Bacillus subtilis* (*Firmicutes*: *Bacilli*) but promote the growth and metabolic activity of *Escherichia coli* (*Gammaproteobacteria*: *Enterobacterales*). Our previous study demonstrated that five-millimolar diaphorin, which exhibits significant inhibitory and promoting effects on cultured *B. subtilis* and *E. coli*, respectively, inhibits *in vitro* gene expression utilizing purified *B. subtilis* and *E. coli* ribosomes. This suggested that the adverse effects of diaphorin on *B. subtilis* are partly due to its influence on gene expression. However, the result appeared inconsistent with the positive effects on *E. coli*. Moreover, the diaphorin concentration in bacterial cells, where genes are expressed *in vivo*, may be lower than in culture media. Therefore, the present study analyzed the effects of 50 and 500 μM of diaphorin on bacterial gene expression using the same analytical method. The result revealed that this concentration range of diaphorin, in contrast to five-millimolar diaphorin, promotes the *in vitro* translation with the *B. subtilis* and *E. coli* ribosomes, suggesting that the positive effects of diaphorin on *E. coli* are due to its direct effects on translation. This study demonstrated for the first time that a pederin-type compound promotes gene expression, establishing a basis for utilizing its potential in pest management and industrial applications.

**Importance:** This study revealed that a limited concentration range of diaphorin, a secondary metabolite produced by a bacterial symbiont of an agricultural pest, promotes cell-free gene expression utilizing substrates and proteins purified from bacteria. The unique property of diaphorin, which is inhibitory to various eukaryotes and *Bacillus subtilis* but promotes the growth and metabolic activity of *Escherichia coli*, may affect the microbial flora of the pest insect, potentially influencing the transmission of devastating plant pathogens. Moreover, the activity may be exploited to improve the efficacy of industrial production by *E. coli*, which is often used to produce various important materials, including pharmaceuticals, enzymes, amino acids, and biofuels. This study elucidated a part of the mechanism by which the unique activity of diaphorin is expressed, constructing a foundation for applying the unique property to pest management and industrial use.

---

Microbes utilize secondary metabolites to mediate interactions with neighboring organisms. Such molecules exhibit diverse biological activities, some of which facilitate symbiotic relationships between the microbes and their animal hosts (1, 2).

Diaphorin is a polyketide produced by “*Candidatus* Profftella armatura” (*Gammaproteobacteria*: *Burkholderiales*), an intracellular symbiont harbored alongside the primary symbiont “*Candidatus* Carsonella ruddii” (*Gammaproteobacteria*: *Oceanospirillales*) (3, 4) in the bacteriome organ (5–7) of the Asian citrus psyllid *Diaphorina citri* (Hemiptera: Psyllidae) (8–11). *D. citri* is a serious agricultural pest that transmits “*Candidatus* Liberibacter” spp. (*Alphaproteobacteria*: *Rhizobiales*), the pathogens of the most destructive and incurable citrus disease, huanglongbing (12, 13). Conserved presence of *Profftella* and its diaphorin-synthesizing gene clusters in *Diaphorina* spp. underline the physiological and ecological significance of diaphorin for the host psyllids (14, 15). Diaphorin, which *D. citri* contains at a concentration as high as 2–20 mM in the body (16), exerts inhibitory effects on various eukaryotes (8, 17, 18) and *Bacillus subtilis* (*Firmicutes*: *Bacilli*) (19) but promotes the growth and metabolic activity of *Escherichia coli* (*Gammaproteobacteria*: *Enterobacterales*) (19), implying that this secondary metabolite serves as a defensive agent of the holobiont (host-symbiont assemblage) against eukaryotes and some bacterial lineages but is beneficial for other bacteria (8, 17, 19). Besides “*Ca*. Liberibacter” spp. and the bacteriome-associated mutualists, *D. citri* may harbor various secondary symbionts of a facultative nature, including *Wolbachia* (*Alphaproteobacteria*: *Rickettsiales*) and *Arsenophonus* (*Gammaproteobacteria*: *Enterobacterales*) (14). Recent studies are revealing that interactions among these bacterial populations are important for psyllid biology and host plant pathology (10, 14, 20–22). In this context, the unique property of diaphorin may affect the microbiota of *D. citri*, potentially influencing the transmission of “*Ca*. Liberibacter” spp. Moreover, this unique activity of diaphorin may be exploited to improve the efficacy of industrial production by *E. coli*, which is frequently used to produce various important materials, including pharmaceuticals, enzymes, amino acids, and biofuels (19).

Diaphorin belongs to the family of pederin-type compounds (8, 19), which exhibit toxicity and antitumor activity by suppressing eukaryotic protein synthesis through binding to the E-site of the 60S subunit of eukaryotic ribosomes (23). However, little is known about the effects of these compounds on bacterial gene expression (24). To explore the possibility that diaphorin exerts its distinct activity on bacteria by directly targeting bacterial gene expression, our previous study analyzed the effects of diaphorin on the *in vitro* gene expression using ribosomes isolated from *B. subtilis* and *E. coli*, quantifying production of the super folder green fluorescent protein (sfGFP) (25). Five-millimolar diaphorin was used for the analysis because this concentration exhibited significant inhibitory and promoting effects on *B. subtilis* and *E. coli*, respectively, in culture experiments (19). The result showed that five-millimolar diaphorin inhibits gene expression involving ribosomes from both *B. subtilis* and *E. coli*, suggesting that the adverse effects of diaphorin on *B. subtilis* are attributed to, at least partly, its inhibitory effects on gene expression (25). On the other hand, the result did not explain the promoting effects of diaphorin on *E. coli*. Moreover, the concentration of diaphorin in the intracellular environment, where the inherent gene expression machinery works, may be lower than in the culture medium. Therefore, in the present study, we analyzed the effect of 50 and 500 μM of diaphorin on bacterial gene expression using the same assay system.

Cell-free translation of sfGFP with diaphorin at final concentrations of 50 and 500 μM demonstrated that this concentration range of diaphorin promotes the *in vitro* gene expression involving ribosomes of both *E. coli* and *B. subtilis* (Fig. 1). Namely, the relative activity of gene expression using the *E. coli* ribosome treated with 50 μM diaphorin was 1.079 ± 0.012 (mean ± standard error, *n* = 48), which was moderately (7.9%) but significantly (*p* < 0.001, Steel test) higher than that of the control (1.000 ± 0.008, *n* = 96, Fig. 1A). Furthermore, the relative gene expression activity using the *E. coli* ribosome treated with 500 μM diaphorin was 1.089 ± 0.017 (*n* = 48), which was again moderately (8.9%) but significantly (*p* < 0.001, Steel test) higher than that of the control (Fig. 1A). These results imply that the positive effects of diaphorin on the growth and metabolic activity of *E. coli* (19) can be attributed to its direct effects on the core gene expression machinery. When cultured in media containing five-millimolar diaphorin (19), *E. coli* may be able to keep the intracellular diaphorin concentration within this range, positively affecting their vital activities. Regarding *B. subtilis*, although the relative gene expression activity using the *B. subtilis* ribosome along with 50 μM diaphorin (0.992 ± 0.023, *n* = 48) was not significantly different (*p* > 0.05, Steel test, Fig. 1B) from the control (1.000 ± 0.011, *n* = 96), the gene expression using the *B. subtilis* ribosome with 500 μM diaphorin (1.084 ± 0.034, *n* = 48) was moderately (8.4%) but significantly (*p* < 0.001, Steel test) higher than the control (Fig. 1B). This result appears inconsistent with previously observed adverse effects of the same concentration of diaphorin on the cultured *B. subtilis* (19). However, transmission electron microscopy showed that diaphorin also damages the *B. subtilis* cell envelope (19), which may negate the positive effects of the appropriate concentration of diaphorin on the gene expression machinery of *B. subtilis*.

**Figure 1.**
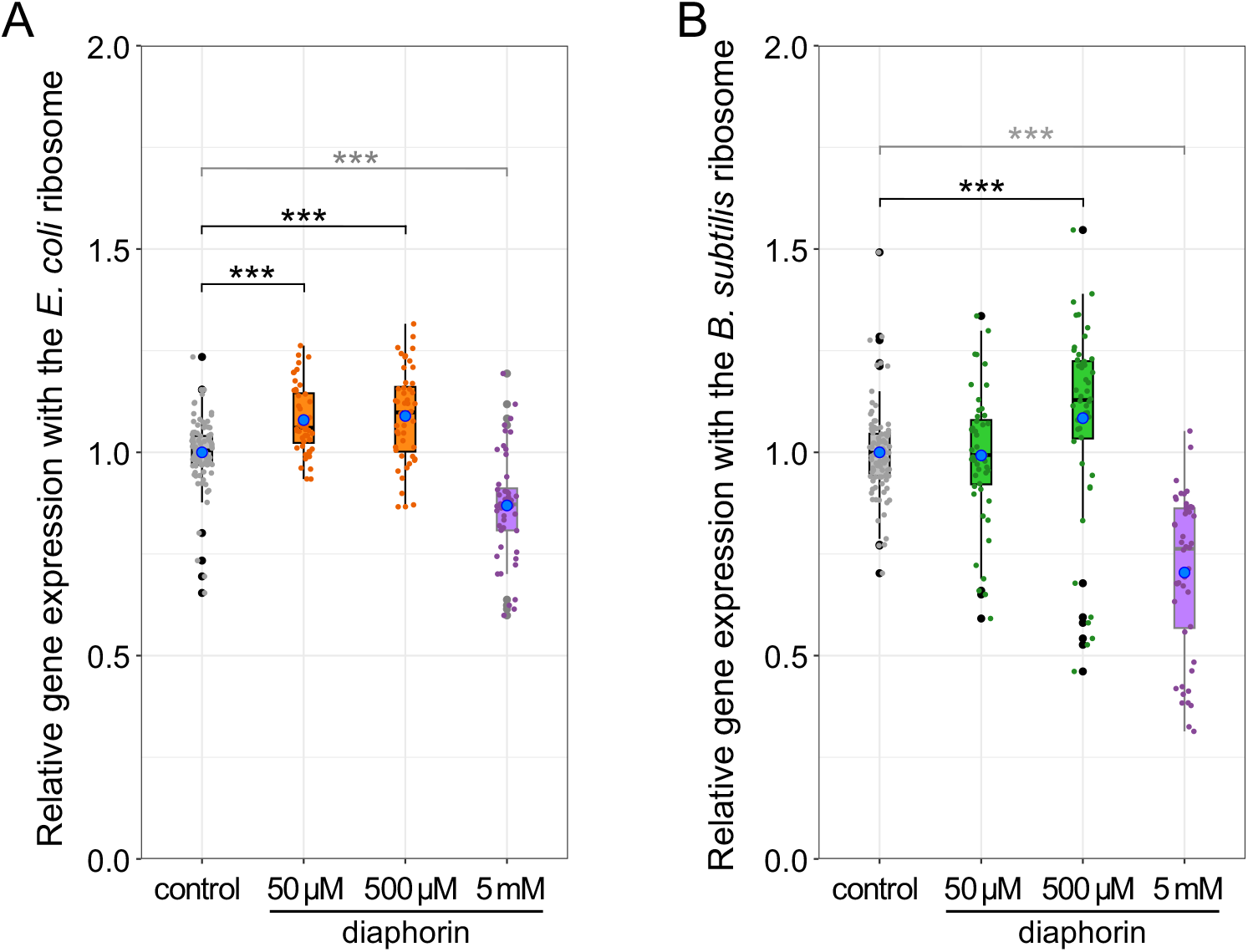
Cell-free gene expression with bacterial ribosomes is promoted by a limited concentration range of diaphorin. (A) Relative gene expression with the *E. coli* ribosome. The signal intensity of synthesized sfGFP in each sample is normalized to the mean signal intensity of control samples. Jitter plots of all data points (control, *n* = 96; others, *n* = 48) and box plots (gray, control; orange, 50 μm and 500 μm diaphorin) showing their distributions (median, quartiles, minimum, and maximum) are indicated. Blue dots represent the mean. Asterisks indicate a statistically significant difference (***, *p* < 0.001, Steel test). For reference, previously published data of 5 mM-diaphorin treatment (19) are shown in purple dots (*n* = 48) with a box plot. (B) Relative gene expression with the *B. subtilis* ribosome. The signal intensity of synthesized sfGFP in each sample is normalized to the mean signal intensity of control samples. Jitter plots of all data points (control, *n* = 96; others, *n* = 48) and box plots (gray, control; green, 50 μm and 500 μm diaphorin) showing their distributions (median, quartiles, minimum, and maximum) are indicated. Blue dots represent the mean. Asterisks indicate a statistically significant difference (***, *p* < 0.001, Steel test). Previously published data of 5 mM-diaphorin treatment (19) are shown in purple dots (*n* = 48) and a box plot.

This study elucidated a part of the mechanism by which the unique activity of diaphorin is expressed, constructing a foundation for applying the unique property of diaphorin to pest management and industrial use. Moreover, this study demonstrated for the first time that a pederin-type compound promotes the gene expression of organisms.

## Materials and methods

### Preparation of diaphorin

Diaphorin was extracted and purified as described previously (8, 17, 19, 25). Adult *D. citri* were ground in methanol, and the extracts were purified using an LC10 high-performance liquid chromatography system (Shimadzu) with an Inertsil ODS-3 C18 reverse-phase preparative column (GL Science).

### Preparation of the *Bacillus subtilis* ribosome

The *B. subtilis* ribosomes were purified as described previously (25). *B. subtilis* cells were passed through a French press cell (Ohtake) at approximately 110 MPa (16,000 psi), and ribosomes were captured using HiTrap Butyl FF columns (Cytiva). The eluent was ultracentrifuged (100,000 ×g, 4°C, 16 h) using Optima L-100 XP Ultracentrifuge (Beckman Coulter) to sediment ribosomes.

### Quantification of cell-free synthesis of sfGFP

The *in vitro* gene expression activities involving ribosomes of *E. coli* and *B. subtilis* were evaluated utilizing a PURE*frex 2*.*0* kit (GeneFrontier) as previously described (25). Reaction solutions of translation were separated by SDS-polyacrylamide gel electrophoresis. After renaturation, the fluorescence of sfGFP was elicited at 488 nm, passed through a 520 nm band pass filter, and recorded using a Typhoon 9400 image analyzer (GE Healthcare). The fluorescence intensity of sfGFP was quantified using the ImageQuant TL software (version 8.1, GE Healthcare).

### Statistical analysis

All statistical analyses were conducted using R version 4.1.3. Multiple comparisons were conducted using the Kruskal-Wallis test followed by the Steel test.

## Acknowledgments

This work was supported by the Japan Society for the Promotion of Science (https://www.jsps.go.jp) KAKENHI (grant number 20H02998) to A.N. The funder had no role in study design, data collection and analysis, decision to publish, or manuscript preparation.

